# Predictive coding during action observation revealed by human electrocorticographic activity

**DOI:** 10.1101/2022.12.13.519734

**Authors:** Chaoyi Qin, Frederic Michon, Yoshiyuki Onuki, Yohei Ishishita, Keisuke Otani, Kensuke Kawai, Pascal Fries, Valeria Gazzola, Christian Keysers

## Abstract

Predictive coding is a theoretical framework that has received much attention for its ability to generate testable hypotheses on how multiple brain regions integrate information during cognitive functions. Given relatively large sensorimotor delays, during social interactions, predicting the behavior of others is crucial to enable joint actions or provide competitive advantages. The action observation network (AON) has been extensively studied, but how information is integrated across its main nodes remains poorly understood. Here we leverage the high spatial and temporal resolution of intracranial Electrocorticography (ECoG), to characterize how the key nodes of the AON - including precentral, supramarginal and visual areas - exchange information. We found more top-down beta oscillation from precentral to supramarginal contacts during the observation of predictable actions while more bottom-up gamma oscillation from visual to supramarginal contacts were measured for unpredictable actions. These results, in line with predictive coding, provide critical evidence towards an understanding of how nodes of the AON integrate information to process the actions of others.

## Introduction

The idea that the brain is a predictive coding machine has received increasing interest(Bastos et al., 2012; Friston, 2005). It has been proposed that the brain operates by generating predictions about upcoming events that are sent in feedback fashion from hierarchically higher to lower regions in the beta-band. These predictions are compared against sensory input, and the difference - the so called prediction error - is then sent in a feedforward fashion from hierarchically lower to higher regions in higher-frequency bands, in particular in the gamma band(Bastos et al., 2012; Fries, 2015; Friston et al., 2015).

A particular domain in which predictive coding merits attention is the domain of action observation. The network of brain regions involved in action observation (AON) is increasingly well mapped. It includes nodes around the medial occipital, supramarginal and precentral gyri that are activated during the execution of similar actions (Caspers et al., 2010; Gazzola and Keysers, 2009; Rizzolatti and Sinigaglia, 2016). However, despite two decades of intense investigation, it remains poorly understood how these nodes interact and integrate information while witnessing sequences of actions. Several papers have suggested that the predictive coding framework may help structure the way we think of the information flow within this system (Friston et al., 2011; Keysers and Gazzola, 2014; Keysers and Perrett, 2004; Kilner and Frith, 2008). Indirect evidence that predictions might be computed within this AON comes from single cell recording in monkeys, in which the response to a particular action can depend on what action can be predicted to come next (Bonini et al., 2010; Umiltà et al., 2001), and from fMRI studies showing that action representations in the parietal and premotor nodes depend on the sequence in which acts are presented(Thomas et al., 2018). Importantly, predictive coding makes a simple, testable prediction: if we observe sequences of actions that allow us to predict what act will come next (e.g. that after grasping a glass and bringing it to your mouth, you are likely to drink from it), we should observe increased feedback predictions and reduced feedforward prediction errors compared to the same actions shown in random - and hence unpredictable - order. More traditional accounts of action observation that simply posit the presence of a hierarchy of brain regions that classify observed actions through feedforward processing alone, do not predict such increased feedback and reduced feedforward information flow for predictable over unpredictable sequences of actions.

Recently, we used depth-resolved 7T fMRI in combination with intersubject correlation analysis to test this account by using the fact that feedback connections have a specific spatial profile(Cerliani et al., 2021): feedback from premotor to inferior parietal regions are known to terminate in layers 3, 5 and 6 in the monkey, and we showed that indeed, intersubject correlation was increased at depths aligning with these layers for predictable actions, and intersubject functional connectivity confirmed that this effect could originate from premotor feedback.

As attributing BOLD activity at certain depths to feedback or feedforward information remains tentative, particularly outside of the visual cortices(Finn et al., 2020), here we aim to leverage the temporal resolution of electrocorticography to shed new light on this conceptually important issue. In particular, there is mounting evidence that the signaling of feedback predictions, and feedforward prediction errors, respectively, are associated with directed information transfer in distinguishable frequency bands, namely the beta and gamma band, respectively(Andre M. Bastos et al., 2015; Bastos et al., 2012; Fries, 2015). Between the premotor and parietal lobes, the high-beta (20-30Hz) and gamma (60-90Hz) frequency bands seem particularly involved in the integration of information (Mooshagian et al., 2021; Tia et al., 2017). If we use the stimuli from our fMRI studies (Table 1), in which participants viewed the same acts in either predictable (intact) sequences or in unpredictable (temporally scrambled) sequences, matched for low-level features using camera changes for both sequences (Figure 1 and Supplementary Figure S1), the predictive coding account makes two testable predictions: the supramarginal gyrus (SMG) should receive more precentral (PreCG) feedback in the high-beta range for intact sequences, and more feedforward prediction errors from the middle occipital gyrus (MOG) for the scrambled sequences (Fig. 1A-B). We thus selected ECoG electrodes in the precentral, supramarginal cortex and middle occipital gyrus across 10 patients implanted with ECoG grids (Fig.1 C). We selected those three regions, because they encompass key nodes of the action observation network, and because we had a sufficient number of patients with ECoG strips that encompassed pairs of these regions to calculate measures of directed information transfer: 7 patients had electrodes in the PreCG and SMG and 6 had electrodes in the SMG and MOG (Fig. 1D).

**Table 1:** Stimuli. List of sequences used as stimuli with total duration in seconds and number of motor acts shown. The first 12 were rated as familiar to Japanese individuals based on an informal evaluation by experimenter YO. The remaining 8 (labeled with letters a-h) from the original study by Thomas et al. (2018) were not used, because they were considered less familiar to Japanese individuals.

|  | **Action** | **Seconds** | **Acts** |
| --- | --- | --- | --- |
| 1 | Inflating and tying a balloon | 51 | 27 |
| 2 | Preparing bread with butter and jam | 79 | 40 |
| 3 | Sewing a button | 66 | 42 |
| 4 | Writing a gift card | 83 | 39 |
| 5 | Arranging flowers in a vase | 82 | 39 |
| 6 | Framing a picture | 112 | 39 |
| 7 | Cleaning spectacles | 69 | 38 |
| 8 | Cleaning a laptop screen | 46 | 28 |
| 9 | Sending a letter | 42 | 34 |
| 10 | Replacing battery in a torch | 51 | 27 |
| 11 | Replacing a pillow cover | 44 | 35 |
| 12 | Folding a shirt | 38 | 20 |
| a | Toasting bread | 65 | 30 |
| b | Making a paper boat | 94 | 32 |
| c | Rolling a cigarette | 72 | 30 |
| d | Applying nail polish | 49 | 23 |
| e | Squeezing oranges | 62 | 40 |
| f | Sharpening a pencil. | 83 | 44 |
| g | Removing nail polish | 64 | 32 |
| h | Preparing a sandwich | 77 | 27 |

**Figure 1:**
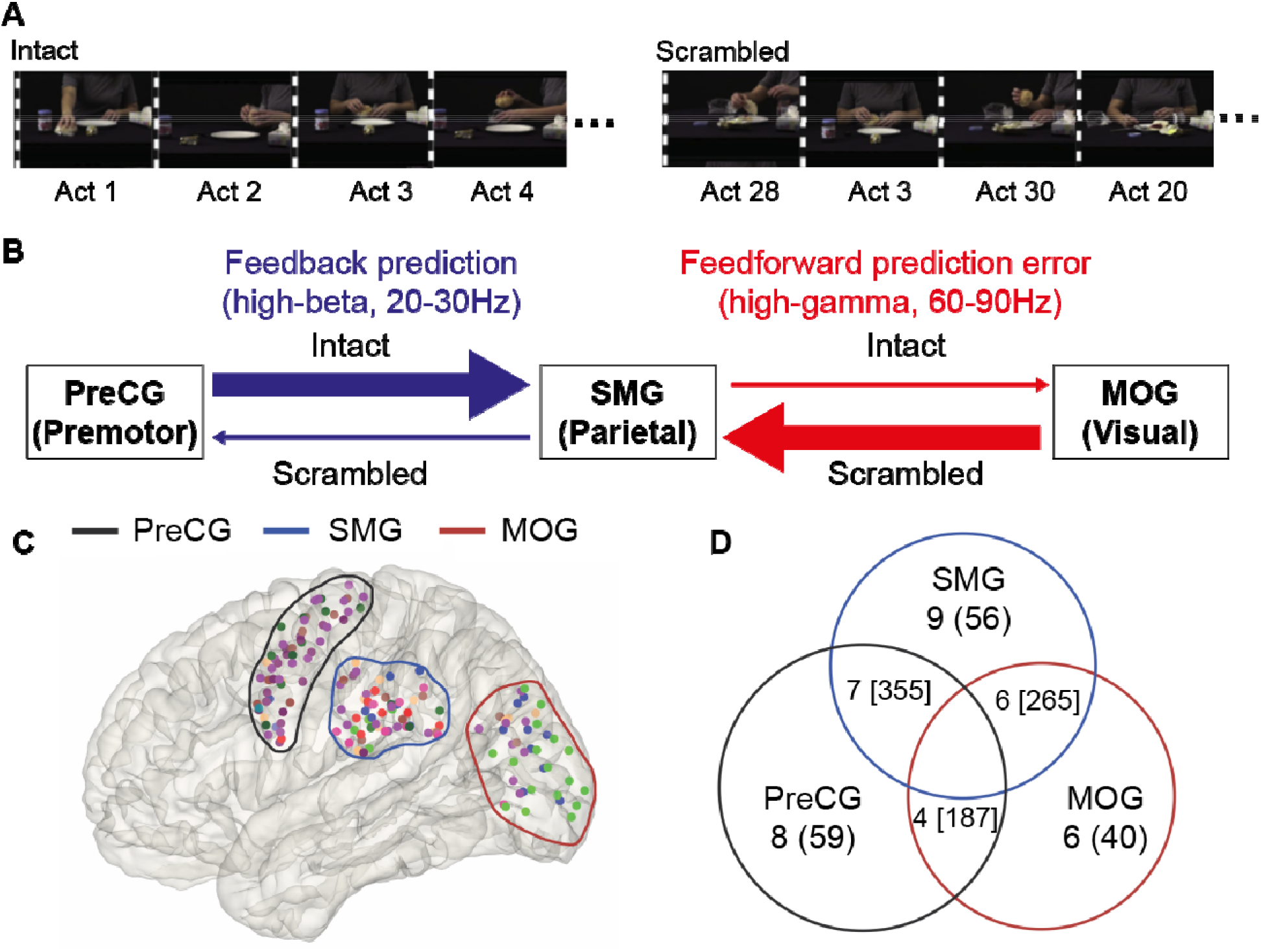
Stimuli and hypotheses. **(A)** We presented participants with movies of everyday hand actions lasting ~1 min in length, filmed simultaneously with two cameras 45° apart, that were cut into ~30 individual motor acts lasting ~2 s (Table 1). In the intact condition, the motor acts were presented in their original order, but switching from one camera-view to the other at the transition between acts. In the scrambled condition, the acts were presented in randomized order. The change between camera-views were introduced in both conditions, because randomizing would otherwise have introduced visual transients in the Scrambled but not in the Intact movie. With these camera changes, intact and scrambled sequences are matched for motion-energy (Supplementary Figure S1). **(B)** Based on predictive coding, we hypothesize that for Intact sequences (top arrows), the parietal node of the action observation network in the supramarginal gyrus (SMG) would receive comparatively more feedback (blue) from the premotor nodes in the precentral gyrus (PreCG) but less feedforward (red) prediction errors from high-level visual cortices in the middle occipital gyrus (MOG), compared to the scrambled condition. We expect feedforward prediction errors to be mainly in the gamma range (60-90Hz), and feedback signals to be mainly in the high-beta range (20-30Hz), the size and direction of the arrows represents the relative strength of coherence and PSI respectively. **(C)** The spatial distribution of electrodes in the three regions on a glass brain in MNI space, the colored circles depict the rough boundaries of these regions. **(D)** The numbers outside the parentheses represent the number of patients who have electrodes in this region or across two regions. The numbers inside the parentheses represent the total number of recording electrodes in this region. The numbers inside the square brackets represent the total number of recording electrode pairs across two regions.

## Results

### Increased high-beta power in precentral and supramarginal channels for predictable actions

To investigate whether beta oscillations were indeed increased for predictable action sequences, as an increase in feedback information may suggest, we calculated the power spectral density (PSD) for all movies in each condition and used a linear mixed effect model (LME) to compare power across conditions. We found that in the high-beta range, precentral and supramarginal regions showed the hypothesized increase in power for the more predictable intact compared to the less predictable scrambled condition (p<0.05, corrected) (Fig. 2A and 2B). When we averaged the power at the hypothesized high-beta frequency range (20-30Hz) in each person across all electrodes, we saw that the majority of subjects had higher beta power in the intact condition, in both the precentral (t_(116)_ = 3.09, p = 0.002) (Fig. 2A inset left) and the supramarginal region (t_(110)_ = 2.29, p = 0.02) (Fig. 2B inset left).

**Figure 2:**
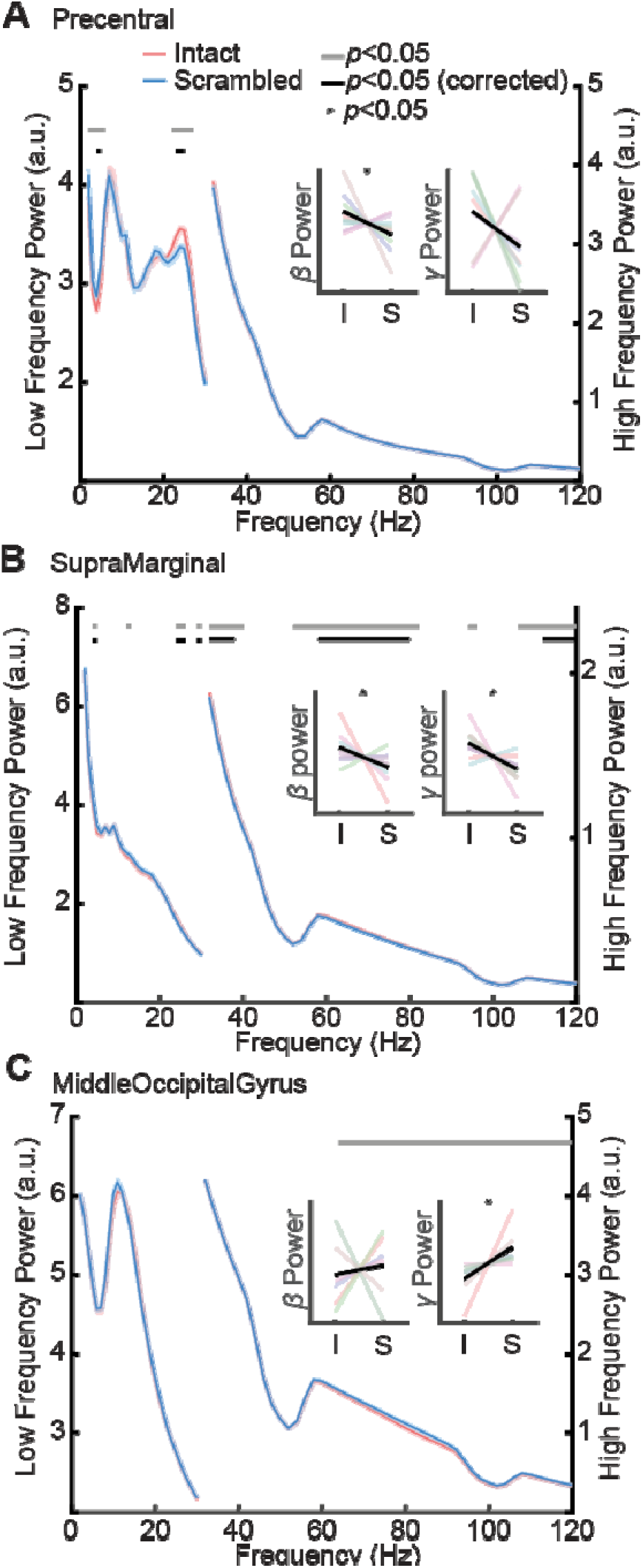
Power spectral density. The power spectral density functions estimated separately for the low and high frequency range across the whole movie viewing period, the insets show the average power of high-beta (20–30Hz) and gamma (60–90Hz) in both Intact (I) and Scrambled (S) conditions for each subject. (A) Precentral channels showed significantly higher power in Intact conditions from 23 to 25 Hz (FDR corrected p < 0.05) and a similar trend was observed in beta power averaged from 20 to 30Hz (Inset on the left). (B) Supramarginal channels showed significantly higher power in Intact condition from 24 to 26, 29 to 30, 58 to 80 and 112 to 120 Hz (FDR corrected p < 0.05), and a similar effect was also observed in beta power averaged from 20 to 30 Hz (inset on the left) as well as gamma power averaged from 60 to 90 Hz (inset on the right). (C) Middle-occipital channels showed significantly higher power in Scrambled conditions from 64 to 120 Hz (uncorrected p < 0.05), and the same effect was observed in gamma power averaged from 60 to 90 Hz (inset on the right).

### Increased gamma power in middle-occipital channels for unpredictable actions

To investigate the notion that unpredictable actions show increased feedforward signals in the gamma range in earlier visual cortices, we quantified the power in the gamma range. We observed opposite effects in earlier visual cortices and supramarginal cortices, with significantly higher gamma power in intact compared to scrambled condition over supramarginal cortices (p<0.05, Fig. 2B) and the hypothesized increase of gamma power for the scrambled movies compared to the intact movies in channels over the middle occipital cortices (p<0.05, Fig. 2C). Examining the averaged power in the gamma band(60-90Hz), we found that most of the participants had higher gamma power in the Intact condition in the supramarginal (t_(110)_ = 3.12, p = 0.002) (Fig. 2B inset right), but higher gamma power in the Scrambled condition in the middle-occipital cortices (t_(78)_ = −2.64, p = 0.01) (Fig. 2C inset right).

### Temporal dynamics of beta and gamma power relative to camera change

We next investigated the temporal dynamics of both high-beta and gamma power in these regions just before and after the camera changes. The camera changes represent a challenging event for the brain. From a low-level visual point of view, camera changes lead to a sudden change in the visual input, similar to those occurring during saccades though not self-initiated like saccades. During saccades, predictions from higher-visual areas are thought to maintain a sense of continuity in the visual scene. We may thus expect an interplay between putatively predictive signals in the high-beta range preparing the system for a camera change ramping up around the likely time of a camera-change, followed by a high-gamma signal starting in the occipital regions caused by the strong change in visual input following the camera change. The former might be more pronounced for intact sequences, and the latter for scrambled sequences.

In the high-beta range, all regions showed a pattern in which power was higher around the time of the camera change than in the middle of a segment. As expected, for precentral and supramarginal channels, the power was higher for the intact sequences, which encourage predictions, at several time-points (p < 0.05, Fig. 3A-C), with significantly higher beta power in the precentral area from 140 ms before the camera change until 220ms after the camera change and in the supramarginal area from 380 until 740 ms after the camera change (Fig. 3A,B). The middle occipital cortex however showed a different pattern, with higher high-beta for intact movies only appearing late in the segment, with early segments showing the opposite pattern (albeing only at uncorrected levels; Figure 3C).

**Figure 3:**
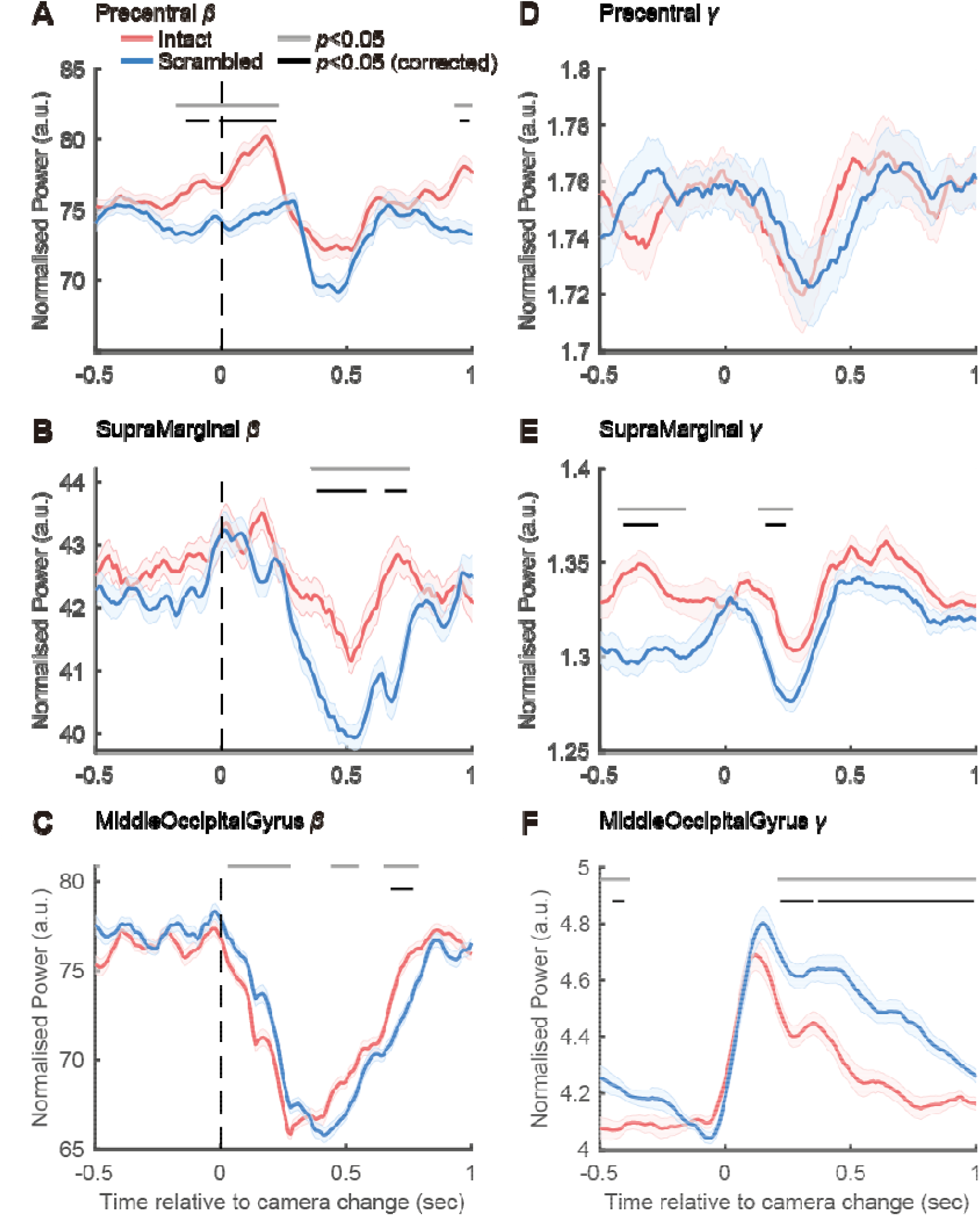
Temporal dynamics of beta and gamma power. Time courses of averaged beta (20–30Hz, left column) and gamma (60–90Hz, right column) power in Intact (red lines) and Scrambled (blue lines) conditions in each region. (A) Precentral showed significantly higher beta power in Intact condition even 140 ms before camera change, which lasted till 220 ms after the camera change. (B) Supramarginal showed significantly higher beta power in Intact condition from 380 ms after the camera change. (C) Middleoccipital showed significantly higher beta power in Intact condition from 680 ms after the camera change. (D) Precentral showed no gamma power difference between conditions. (E) Supramarginal showed significantly higher gamma power in Intact condition from 410 ms before and 160 ms after the camera change. (F) Middleoccipital showed significantly higher gamma power in Scrambled condition from 220 ms after the camera change.

With regard to high-gamma power, the middle occipital cortex showed the expected increase in high-gamma after typical visual latencies, with the power being higher for the scrambled sequences that should generate the highest feedforward prediction errors (Fig. 3F). The supramarginal and precentral cortices failed to show such a high-gamma peak following the sudden change in visual input, and instead showed a dip in high-gamma power, which in the supramarginal cortex was more pronounced for the scrambled sequences (Fig. 3E). Precentral channels, on the other hand, showed no such changes in gamma power (Fig. 3D).

### Beta synchronization and information transfer between precentral and supramarginal

The increased high-beta activities in precentral channels for intact sequences preceded that in supramarginal channels. This would be in line with a model in which predictions in premotor regions would be transferred backwards to the parietal nodes of the action observation network. To test this notion, we computed the interregional connectivity across precentral and supramarginal channels using spectral coherence.

First, we measured imaginary coherence (see Methods) using all electrode pairs within the first second after the camera change, and found significantly higher beta coherence between precentral and supramarginal channels in the intact compared to the scrambled condition. This effect was restricted to the high-beta range (23-30Hz, *p*_corr_<0.05) (Fig. 4A). Interestingly, the low-beta range showed a difference in the opposite direction, confirming the functional dissociation between low- and high-beta. Frequencies around 50Hz were masked out due to line noise contaminating the coherence estimates. Furthermore, using a sliding window method to characterize the timing of the differential high-beta coherence, we find that it emerges 300ms after the camera change (p<0.05) (Fig. 4C).

**Figure 4:**
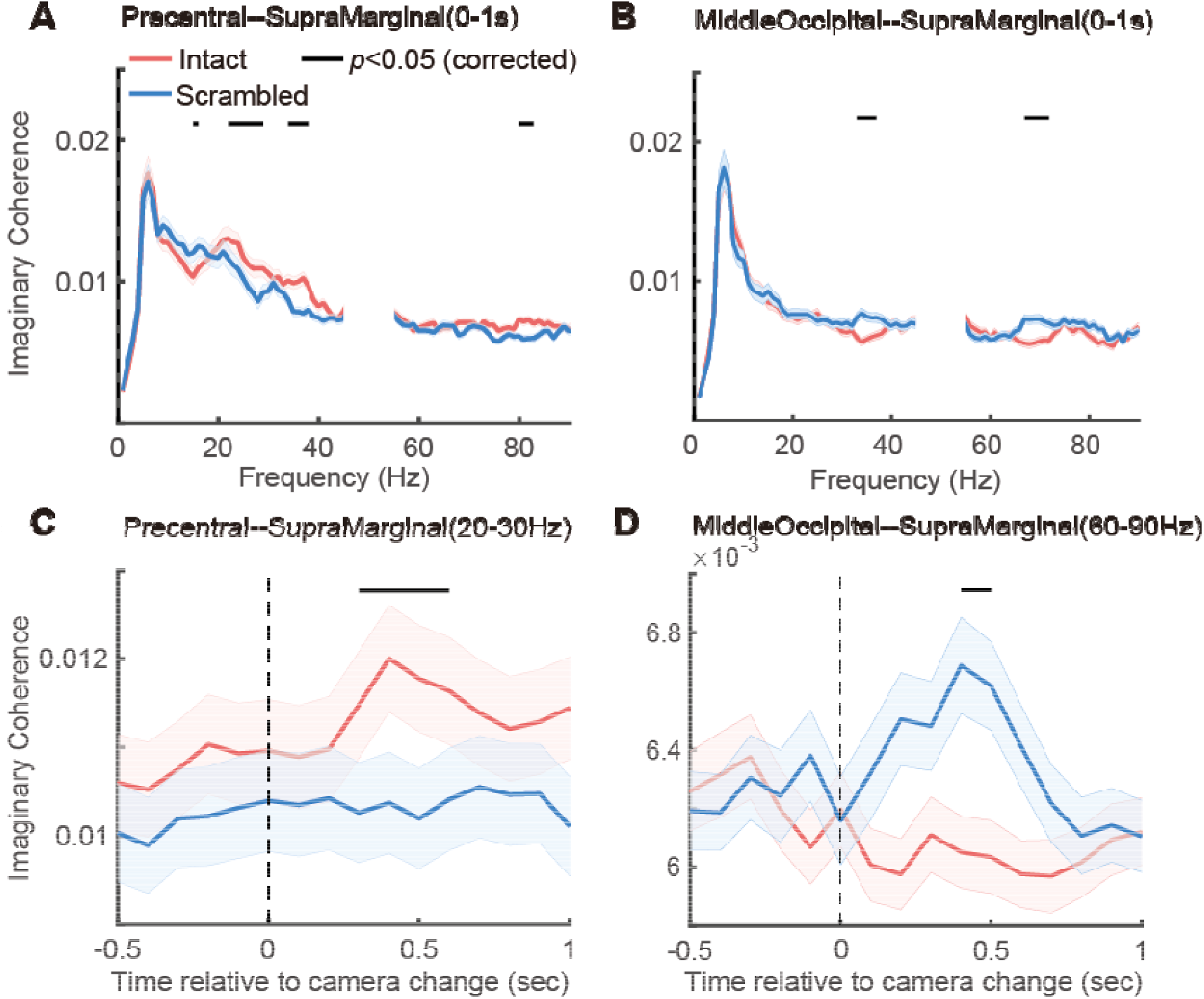
Spectral Coherence and temporal dynamics of coherence. (A) Precentral and supramarginal showed significantly higher coherence in intact condition in beta frequency from 23 to 30 Hz (FDR corrected p < 0.05). (B) Middleoccipital and supramarginal showed significantly higher coherence in scrambled condition in gamma frequency from 68 to 73 Hz (FDR corrected p < 0.05). (C) Precentral and supramarginal showed significantly higher coherence in intact condition from 300ms after camera change (FDR corrected p < 0.05). (D) Middleoccipital and supramarginal showed significantly higher coherence in scrambled condition from 400ms after camera change (FDR corrected p < 0.05).

To further investigate the directionality of information transfer between the precentral and supramarginal, we calculated the non-parametric Granger Causality (GC) using all electrode pairs within the first second after the camera change. In the intact compared to the scrambled condition, the GC spectrum exhibited higher feedback information from precentral to supramarginal in the high-beta band (p<0.05; Fig. 5A), but there was no difference between conditions in the opposite (feedforward) direction (Fig. 5B). Both GC and coherence exhibited frequencies that were in the high beta range (20-30Hz). We estimated the Phase Slope Index (PSI) in 20-30 Hz using a sliding window method to further validate the information flow between precentral and supramarginal in this high-beta range. Starting 400ms before the camera change, the PSI exhibited opposing information directions, with feedforward information from supramarginal to precentral in scrambled state and feedback information from precentral to supramarginal in intact condition (Fig. 6A).

**Figure 5:**
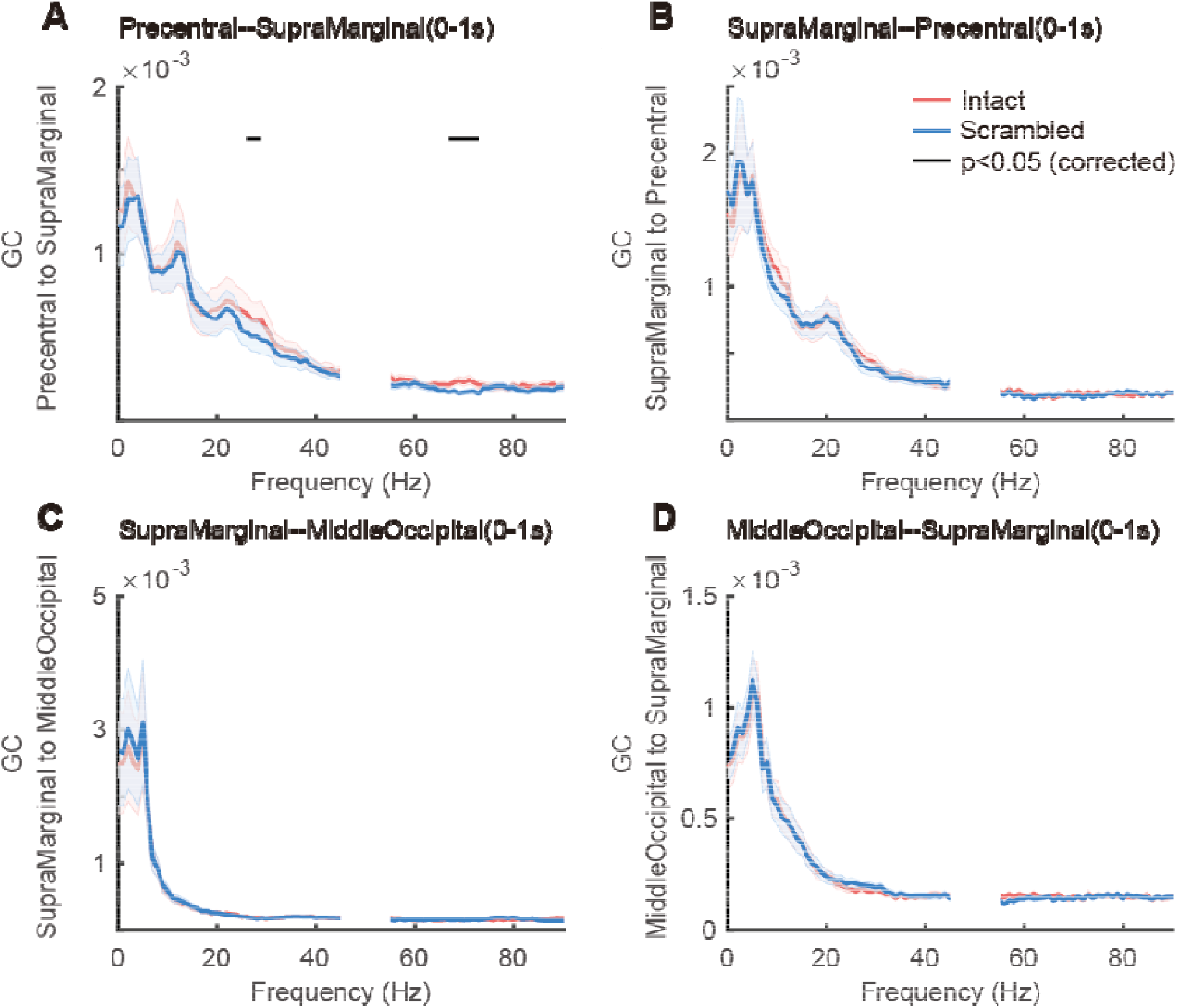
Non-parametric Granger Causality. (A) Stronger granger causality was found from precentral towards supramarginal in intact condition in beta frequency from 26 to 29 Hz. (B) No difference in the evidence was found from supramarginal to precentral between conditions. (CD) No difference in the evidence was found between middle occipital and supramarginal between conditions.

**Figure 6:**
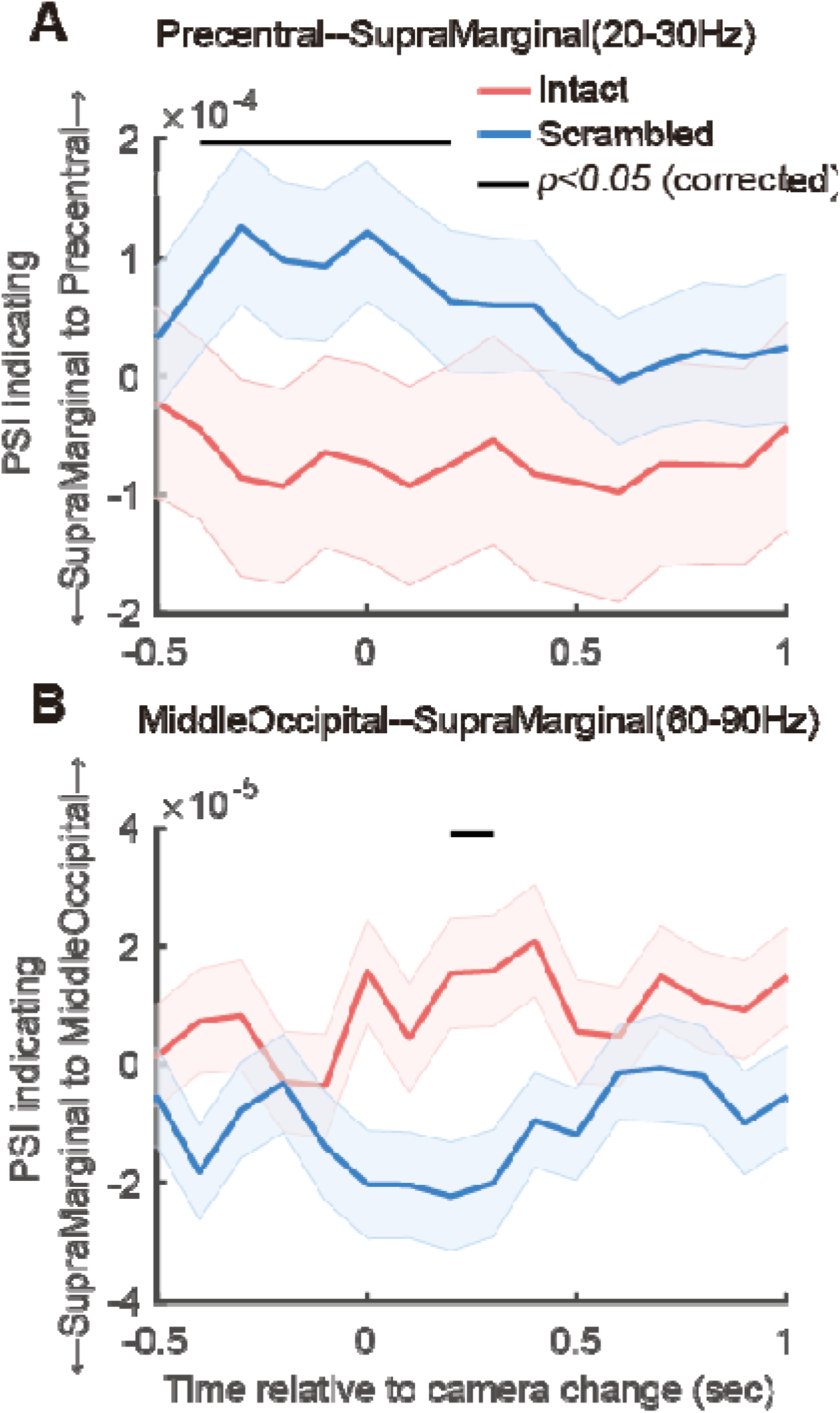
Phase Slope Index. (A) Phase slope index revealed more beta information (20-30Hz) from precentral to supramarginal in intact condition but from supramarginal to precentral in scrambled condition. (B) On the other hand, gamma information (60-90Hz) was observed to transfer more from middleoccipital to supramarginal in scrambled condition but from supramarginal to middleoccipital in intact condition.

### Gamma synchronization and information transfer between supramarginal and middle occipital cortices

We also looked at the interregional relationships between the supramarginal and middle occipital channels. In both the high gamma (60-90Hz) and low gamma (30-40Hz) ranges, the computed imaginary coherence during the first second following the camera change indicated significantly stronger gamma coherence between supramarginal and middle occipital channels in the scrambled condition (p < 0.05, Fig. 4B). Furthermore, the time-resolved coherence increased 400ms after the camera changed in scrambled condition (p < 0.05, Fig. 4D).

Comparing the two conditions using Granger Causality did not demonstrate a preferential information direction between supramarginal and middle occipital channels in the gamma band (at p<0.05, Fig. 5C and D). However, the PSI derived at a high gamma frequency (60-90Hz) suggested the expected direction of effect, with greater feedforward information from the middle occipital to the supramarginal channel, beginning 200ms after the camera change (p<0.05), compared to the intact condition. (Fig. 6B).

## Discussion

How information is integrated and exchanged across the nodes of the action observation network remains poorly understood, but it has been suggested that a predictive coding framework may account for the dominant directions of information flow subserving action observation across this system(Friston et al., 2011; Keysers and Gazzola, 2014; Keysers and Perrett, 2004; Kilner and Frith, 2008). Our findings support this proposal by showing that signals in the high-beta and high-gamma range across the precentral, supramarginal, and middle occipital regions of the AON are differentially modulated by the nature of the action sequences we observe. In what follows, we will discuss changes in these two frequency bands separately, together with some background on how changes in these frequency bands have been associated with feedback and feedforward information flow in the literature.

Beta oscillations in high-level brain regions have been associated with top-down processing and sensori-motor integration via feedback information flow across distal brain regions (Barone and Rossiter, 2021; Andre M. Bastos et al., 2015; André Moraes Bastos et al., 2015; Engel and Fries, 2010; Fries, 2015). Several studies have shown that the power in the beta range, particularly in the high-beta range from 20-30Hz, is modulated by action observation confirming that it may be important for feedback processes also during action observation (Babiloni et al., 2016; Moreno et al., 2013; Muthukumaraswamy and Johnson, 2004; Simon and Mukamel, 2016). Here we found that observation of actions in predictable order caused higher high-beta power in, and higher high-beta coherence between, precentral and supramarginal cortices. While the fact that this power and coherence increase occurs in the beta range is indicative of a feedback direction of information flow, the high temporal resolution of ECoG, and its relatively higher spatial resolution compared to scalp recordings, allows us to directly test the prevalent direction of information flow using phase-slope indices and Granger causality. Both methods confirmed that intact sequences lead to more information flow in the high-beta range in the feedback direction from precentral to supramarginal cortices. This provides what is to our understanding the most direct evidence that predictability of action sequences indeed increases feedback information, in line with the influential but largely untested notion of predictive coding during action observation(Friston et al., 2011; Keysers and Gazzola, 2014; Keysers and Perrett, 2004; Kilner and Frith, 2008), and in line with our findings of increased action-observation related activity for predictable actions in layers of the supramarginal gyrus known to receive premotor feedback(Cerliani et al., 2021). These effects were observed in the high-beta range (above 20Hz), and not in the lower beta range (15-20Hz). The specific role of subbands in the beta range remains poorly understood, but a small number of studies point towards a particular relevance of high-beta for integration of motor signals (Mooshagian et al., 2021; Tia et al., 2017), with the high-beta range selectively altered during the observation of other people’s actions(Simon and Mukamel, 2016).

Gamma power, in contrast, has been associated with local processing (Khanna and Carmena, 2015) and feedforward information flow (Aggarwal et al., 2022; Babiloni et al., 2016; Fries, 2015; van Kerkoerle et al., 2014). In the visual system, gamma is triggered particularly by sudden changes in the visual input (Bartoli et al., 2020; Brunet et al., 2015). In accordance with that literature, the camera changes, which trigger a sudden change in visual input, triggered a transient in high-gamma power in our more classically visual channels in the MOG, and, perhaps unsurprisingly, such camera-change-locked increases were not observed in the SMG and PreCG, where single cell recordings in monkeys have shown many neurons to generalize their responses over changes in low level features and viewpoint(Caggiano et al., 2016; Maranesi et al., 2017) and continue to respond even when critical aspects of the action are occluded(Umiltà et al., 2001). More interestingly, with regard to our predictive coding hypothesis, in the scrambled condition, occipital channels had increased high-gamma power compared to the intact condition, while the supramarginal channels had the reverse tendency. Considering the above mentioned association of high-gamma activity with feedforward(Aggarwal et al., 2022; Richter et al., 2017; van Kerkoerle et al., 2014) and local information(Kopell et al., 2000; Ray and Maunsell, 2011), this increased high-gamma power in the middle occipital cortices could reflect the prediction error that predictive coding models would expect to be generated at a camera change when the visual input does not match what the preceding action would suggest to happen next, as would be the case in our scrambled movies. Importantly, we also found the coherence to be increased in the scrambled condition across the middle occipital cortex and the supramarginal gyrus, and the phase-slope index confirms that this information in the high-gamma band indeed flowed from the middle-occipital to the supramarginal gyrus, in accordance with a predictive coding model. Interestingly, high-gamma activity in the supramarginal gyrus was increased in the intact, compared to the scrambled condition. Given that broadband high-gamma power is known to be tightly linked to neural spiking and thereby reflects local processing(Ray and Maunsell, 2011), this increased high-gamma power in the supramarginal cortex during intact sequences, together with the increased intersubject correlation for intact sequences in the supramarginal gyrus in fMRI BOLD signals(Thomas et al., 2018), suggest that the parietal node indeed represents more than the individual motor acts that are identical across the intact and scrambled movies, and preferentially encodes actions when they integrate into larger, meaningful sequences. Such preferential encoding of longer chains of actions in the parietal node contrasts with the prediction errors that appear to dominate the occipital node, in line with the notion that there is an progressive increase in the ‘temporal receptive field’ of cortical regions along a hierarchy from earlier sensory regions, with activity that integrates information over short intervals to more anterior parietal and frontal regions that can integrate information over minutes(Lerner et al., 2011; Thomas et al., 2018).

After decades of mapping the action observation network using neuroimaging techniques, our electrocorticography data provide unique insights into the poorly understood question of how information is integrated around the well-mapped nodes recruited during action observation. Leveraging the temporal resolution of electrical recordings, and the spatial specificity afforded by recording so close from the cortex, our data shows that when we observe actions in larger, meaningful sequences, when predictions are possible (in our intact sequences), feedback information is transmitted in high-beta oscillation from precentral to supramarginal cortices, and local processing in the high-gamma band is increased in the supramarginal cortices and reduced in occipital visual cortices. This finding is in line with the findings in monkeys, in which neural activity while witnessing predictable actions has shorter latencies in premotor than parietal nodes of the mirror neuron system (Ferroni et al., 2021). When expectations are violated, as following a camera change in our scrambled sequences, a transient increase in high-gamma activity in the visual cortices is triggered, and information flows from these visual cortices to the supramarginal gyrus in the gamma band. Zooming in on the moment of a camera change, our data suggests a particular succession of events. For intact sequences, precentral cortices generate high-beta activity thought to reflect predictions just before the camera change, and this increased high-beta activity continues after the camera change. This pattern is reminiscent of the predictive activity around a saccade-onset in frontal eye fields that is thought to provide the brain with a continuity of perception despite the low-level discontinuity that a saccade causes (Rao et al., 2016; Zirnsak and Moore, 2014). In contrast, when predictions are violated, after a camera change in the scrambled sequences, a high-gamma transient, occurring around the typical response latency of visual areas after the camera change, appears to produce an error signal that is forwarded to the parietal node, where it disrupts local processing (i.e. causes a relative reduction of gamma activity).

Our study has a number of limitations that should be considered. First, here, we focused on three key regions within the AON, while the action observation process is known to be more complex, also involving for instance somatosensory, cerebellar, and other subcortical regions(Abdelgabar et al., 2019; Caspers et al., 2010; Gazzola and Keysers, 2009; Thomas et al., 2018). Future studies involving patients with a wider coverage may be ideally suited to investigate how information from these other nodes may integrate with those of the network we focus on. Second, to increase the statistical power of our analyses, we pooled electrodes over relatively large regions of the cortex within our three regions of interest. Future studies may wish to explore whether specific subregions show different patterns. Indeed, some studies mentioned a specific topological organization of the premotor cortex and its connectivity with the parietal lobe in action observation (Stadler et al., 2012; Urgen and Saygin, 2020). A preliminary analysis of our data however failed to reveal a specific topography of connectivity (Fig S2). Thirdly, while for the high-beta analysis, two methods estimating the direction of information flow (phase-slope index and Granger causality) agree, for the high-gamma frequency range across the occipital and parietal channels, the phase slope index analysis revealed a significant difference between conditions, while the Granger causality did not. The exact source of this discrepancy is difficult to assess,it is suggested that PSI shows relatively more robust performance than Granger causality but such discrepancy temper the confidence we can have in the directionality of the information transfer in the gamma-band across these regions(Bastos and Schoffelen, 2016; Brovelli et al., 2004; Young et al., 2017; Ziehe et al., 2010).

## Methods

### Patients

Ten subjects with refractory epilepsy participated in this study (five males and five females, aged 18-39 years, mean = 27.3 years, standard deviation = 7.3; see Table S1 for the demographic features of patients). Subdural electrodes were placed to localize epileptic foci and examine the cognitive and motor functions of areas under the electrodes. Written informed consent was obtained from all patients. This study was approved by the ethics committee at Jichi Medical University Hospital and registered in the UMIN Clinical Trial Registry (number UMIN000040073).

### Stimuli and experiment procedure

The stimuli used here were a subset of those used in (Thomas et al., 2018). Briefly, twenty movies containing different daily actions (e.g. preparing sandwiches with butter and jam; see Table 1 for the full list) were simultaneously recorded by two video cameras (Sony MC50, 29 frames/s) at an angle of 45 degrees. The videos were edited using Adobe Premiere ProCS5 running on Windows. Each movie was subdivided into shots containing one meaningful motor act each (e.g. taking bread, opening the butter dish, scooping butter with a knife, etc.). This was done on recordings from both camera angles. These motor acts (mean/standard deviation duration 2s ± 1s) were then assembled to build two types of ~1 minute long stimuli (average 67s, Fig. 1). For the Intact (I) presentation, the natural temporal sequence in which the acts were recorded was maintained, but a camera angle change was introduced between every two consecutive acts by alternate sampling from the recordings of the two cameras. In the Scrambled (S) versions, the acts remained the same, but the order of the acts was randomly re-arranged, and a camera angle change was introduced between every two consecutive acts. Camera angle changes were imposed at each act transition in both types of movies to compensate for the visual transients that would otherwise be present only in the scrambled movies. Because that stimulus set depicted actions typical for western europeans, but the experiment was performed in Japan, author YO examined all twenty movies and selected 12 actions that should be familiar to Japanese participants (Table 1). This resulted in 12 intact and 12 scrambled movies. Each movie was presented twice. The experiment was conducted in 6 sessions. They were composed of 3 unique sessions presented twice, with each of the sessions including 4 intact and 4 scrambled movies presented in pseudorandom order, with an inter-movie interval between 8 and 12 s. No behavioral response was required during the experiment, but participants were to carefully observe the videos. Among the included participants, one completed only 3 sessions and one completed only 2 sessions.

### Electrophysiological Recordings and Signal Preprocessing

Intracranial EEG signals were recorded using a Nihon-Kohden system with 1000Hz sampling rate in Jichi Medical University Hospital, Japan. All signals were online referenced to two electrodes in the first head stage. All data analysis was conducted in MATLAB using the fieldtrip toolbox (www.fieldtriptoolbox.org/) and customized scripts. The recorded signals were first low-pass filtered using a 4th order Butterworth filter with a cutoff frequency at 200 Hz. The 50Hz power line noise and its harmonics were removed using bandstop filters with variable bandwidth according to individual power spectra. Channels with obvious artifacts were excluded from further analysis. Each electrode was then locally re-referenced to the average of its neighboring electrodes within 12mm spatial distance. This procedure removes the common recording reference, which otherwise leads to spurious correlations and coherence. Coherence, imaginary coherence, phase-slope index and Granger causality were exclusively calculated between electrodes, for which the neighboring electrodes used for re-referencing had no overlap. Data were down-sampled to 500 Hz for subsequent analyses.

### Electrode Locations and Region Definition

The spatial locations were derived from each patient’s pre-implantation MR images and post-implantation CT images. For each patient, the post-implantation CT was co-registered to the pre-implantation MRI using a six-parameter rigid body transformation, implemented in SPM12 (https://www.fil.ion.ucl.ac.uk/spm/software/spm12/). The registration was visually verified and manually adjusted if necessary. ECoG electrodes were identified semi automatically according to anatomical landmarks in native space (Qin et al., 2017). For visualization of all subjects’ electrodes on an average surface, individual electrode coordinates were transformed to MNI152 space using the Freesurfer CVS function(v6.0.0, surfer.nmr.mgh.harvard.edu/). The regions of interest including precentral gyrus (PreCG), supramarginal gyrus (SMG) and middle occipital gyrus (MOG) were extracted from the Anatomy toolbox(Amunts et al., 2007)(Fig. 1C).

### Trial Separation

In the connectivity analysis including coherence, phase slope index and granger causality, data was first separated into trials based on the time of the camera change. The actions used in our stimuli were composed of a sequence of different shorter acts (e.g. the action of buttering bread included the acts of e.g. taking bread, opening the butter dish, scooping butter with a knife, etc.). These acts were used as “action primitives” and determined the location of the camera changes. The intervals between two camera changes therefore depended by the duration of each act, and varied from 0.4 -- 6.76s. In our analyses, to minimize the overlap between trials as well as preserving the temporal dynamics during action perception, we thus chose a time window from −0.5s to 1s relative to the camera change

### Power Spectral Density

Power spectral density was estimated separately for low(2-30Hz) and high frequencies (30-120Hz). For the low frequencies, the power spectrum was estimated by a short-time fourier transform with Hann tapers of 1s, sliding over the whole experimental session in steps of 0.1s and averaged over all windows in a given condition. For the high-frequency part, we used a multi-taper spectral estimation with 5 tapers in 0.5s windows sliding in steps of 0.1s, and results were also averaged overall time windows in a given condition. Power differences between conditions were then compared using all the electrodes in each selected region of interest.

### Coherence

For the selected time window, all trial data in each electrode within this window were Fourier transformed using multi-tapering with 6Hz frequency smoothing in frequencies ranging from 2 to 120Hz with 1Hz step.The coherence between two signals was then calculated in each electrode pair and each region pair in both conditions, the imaginary part of coherence was taken as the metric to measure the synchronization between regions. The time-resolved coherence was calculated in a sliding-window manner with window length of 1 s and steps of 0.1s and averaged across the frequency points in each frequency band of interest.

### Phase Slope Index

The phase slope index (PSI) was calculated in a similar manner as coherence across regions (Nolte et al., 2008). All trial data for each electrode were Fourier transformed using multi-tapering with 4Hz frequency smoothing and 1Hz frequency resolution. The phase slope index was then calculated across frequencies ranging from 20 to 30Hz and 60 to 90Hz with the same bandwidth of 4Hz for each frequency; the PSI values were then averaged, separately for each of the two frequency bands, to get the temporal dynamics, this was done in each electrode pair for each region pair in both conditions as well.

### Granger Causality

Granger causality (GC) was calculated in a time window from 0–1s relative to the camera change using the nonparametric estimation (Brovelli et al., 2004; Dhamala et al., 2008). The Fourier spectrum was estimated using the same parameters as the PSI mentioned above and entered into a nonparametric spectral matrix factorization as implemented in the FieldTrip toolbox (Oostenveld et al., 2011).

### Statistical Assessment

Statistical assessments were performed to compare the difference between conditions using the LME model implemented in MATLAB. We implemented the LME model with patient and electrode (or electrode pairs) as two random effects and used the restricted maximum likelihood method to optimize. In the model, fixed and random effects were considered together. Post hoc tests of *p* values were performed using FDR correction to correct for multiple comparisons. Statistical inferences were under a significance threshold of p < 0.05 if not specified otherwise.

## Acknowledgements

We thank Teresa de Sanctis for help with preparing the stimuli, and Rajat Mani Thomas for help with data analysis.

## Competing Interests

The authors report no competing interests.

## Funding

Funding: CK was funded by VICI grant 453-15-009, VG by VIDI grant 452-14-015.

## Author Contributions

CK and VG conceived the study, obtained funding, and supervised the analysis of the data. YO acquired the data. YI, KO, KK performed the surgeries. CQ and FM analyzed the data with supervision and advice from PF. CQ and CK wrote the first draft of the paper. All authors edited the manuscript.

## Notes

### Competing Interest Statement

The authors have declared no competing interest.

### Summary of Updates

Add supplementary figure S1 to clarify the difference in the motion energy (low level visual property) of the movie in two conditions.

## References

Abdelgabar AR, Suttrup J, Broersen R, Bhandari R, Picard S, Keysers C, De Zeeuw CI, Gazzola V. 2019. Action perception recruits the cerebellum and is impaired in patients with spinocerebellar ataxia. Brain 142:3791–3805. doi:10.1093/brain/awz337

Aggarwal A, Brennan C, Luo J, Chung H, Contreras D, Kelz MB, Proekt A. 2022. Visual evoked feedforward-feedback traveling waves organize neural activity across the cortical hierarchy in mice. Nat Commun 13:4754. doi:10.1038/s41467-022-32378-x

Amunts K, Schleicher A, Zilles K. 2007. Cytoarchitecture of the cerebral cortex—More than localization. NeuroImage 37:1061–1065. doi:10.1016/j.neuroimage.2007.02.037

Babiloni C, Del Percio C, Vecchio F, Sebastiano F, Di Gennaro G, Quarato PP, Morace R, Pavone L, Soricelli A, Noce G, Esposito V, Rossini PM, Gallese V, Mirabella G. 2016. Alpha, beta and gamma electrocorticographic rhythms in somatosensory, motor, premotor and prefrontal cortical areas differ in movement execution and observation in humans. Clin Neurophysiol Off J Int Fed Clin Neurophysiol 127:641–654. doi:10.1016/j.clinph.2015.04.068

Barone J, Rossiter HE. 2021. Understanding the Role of Sensorimotor Beta Oscillations. Front Syst Neurosci 15:655886. doi:10.3389/fnsys.2021.655886

Bartoli E, Bosking W, Foster BL. 2020. Seeing Visual Gamma Oscillations in a New Light. Trends Cogn Sci 24:501–503. doi:10.1016/j.tics.2020.03.009

Bastos AM, Schoffelen J-M. 2016. A Tutorial Review of Functional Connectivity Analysis Methods and Their Interpretational Pitfalls. Front Syst Neurosci 9. doi:10.3389/fnsys.2015.00175

Bastos AM, Usrey WM, Adams RA, Mangun GR, Fries P, Friston KJ. 2012. Canonical microcircuits for predictive coding. Neuron 76:695–711.

Bastos André Moraes, Vezoli J, Bosman CA, Schoffelen J-M, Oostenveld R, Dowdall JR, De Weerd P, Kennedy H, Fries P. 2015. Visual Areas Exert Feedforward and Feedback Influences through Distinct Frequency Channels. Neuron 85:390–401. doi:10.1016/j.neuron.2014.12.018

Bastos Andre M., Vezoli J, Fries P. 2015. Communication through coherence with inter-areal delays. Curr Opin Neurobiol 31:173–180. doi:10.1016/j.conb.2014.11.001

Bonini L, Rozzi S, Serventi FU, Simone L, Ferrari PF, Fogassi L. 2010. Ventral premotor and inferior parietal cortices make distinct contribution to action organization and intention understanding. Cereb Cortex N Y N 1991 20:1372–1385. doi:10.1093/cercor/bhp200

Brovelli A, Ding M, Ledberg A, Chen Y, Nakamura R, Bressler SL. 2004. Beta oscillations in a large-scale sensorimotor cortical network: Directional influences revealed by Granger causality. Proc Natl Acad Sci 101:9849–9854. doi:10.1073/pnas.0308538101

Brunet N, Bosman CA, Roberts M, Oostenveld R, Womelsdorf T, De Weerd P, Fries P. 2015. Visual Cortical Gamma-Band Activity During Free Viewing of Natural Images. Cereb Cortex 25:918–926. doi:10.1093/cercor/bht280

Caggiano V, Fleischer F, Pomper JK, Giese MA, Thier P. 2016. Mirror Neurons in Monkey Premotor Area F5 Show Tuning for Critical Features of Visual Causality Perception. Curr Biol CB 26:3077–3082. doi:10.1016/j.cub.2016.10.007

Caspers S, Zilles K, Laird AR, Eickhoff SB. 2010. ALE meta-analysis of action observation and imitation in the human brain. NeuroImage 50:1148–1167. doi:10.1016/j.neuroimage.2009.12.112

Cerliani L, Bhandari R, De Angelis L, van der Zwaag W, Bazin P-L, Gazzola V, Keysers C. 2021. Predictive coding during action observation - a depth-resolved intersubject functional correlation study at 7T. bioRxiv 2021.08.30.458143. doi:10.1101/2021.08.30.458143

Dhamala M, Rangarajan G, Ding M. 2008. Analyzing information flow in brain networks with nonparametric Granger causality. NeuroImage 41:354–362. doi:10.1016/j.neuroimage.2008.02.020

Engel AK, Fries P. 2010. Beta-band oscillations—signalling the status quo? Curr Opin Neurobiol 20:156–165. doi:10.1016/j.conb.2010.02.015

Ferroni CG, Albertini D, Lanzilotto M, Livi A, Maranesi M, Bonini L. 2021. Local and system mechanisms for action execution and observation in parietal and premotor cortices. Curr Biol CB 31:2819–2830.e4. doi:10.1016/j.cub.2021.04.034

Finn ES, Huber L, Bandettini PA. 2020. Higher and deeper: Bringing layer fMRI to association cortex. Prog Neurobiol 101930. doi:10.1016/j.pneurobio.2020.101930

Fries P. 2015. Rhythms for Cognition: Communication through Coherence. Neuron 88:220–235. doi:10.1016/j.neuron.2015.09.034

Friston K. 2005. A theory of cortical responses. Philos Trans R Soc Lond B Biol Sci 360:815–836. doi:10.1098/rstb.2005.1622

Friston K, Mattout J, Kilner J. 2011. Action understanding and active inference. Biol Cybern 104:137–160. doi:10.1007/s00422-011-0424-z

Friston KJ, Bastos AM, Pinotsis D, Litvak V. 2015. LFP and oscillations-what do they tell us? Curr Opin Neurobiol 31:1–6. doi:10.1016/j.conb.2014.05.004

Gazzola V, Keysers C. 2009. The Observation and Execution of Actions Share Motor and Somatosensory Voxels in all Tested Subjects: Single-Subject Analyses of Unsmoothed fMRI Data. Cereb Cortex 19:1239–1255. doi:10.1093/cercor/bhn181

Keysers C, Gazzola V. 2014. Hebbian learning and predictive mirror neurons for actions, sensations and emotions. Philos Trans R Soc B Biol Sci 369:20130175. doi:10.1098/rstb.2013.0175

Keysers C, Perrett DI. 2004. Demystifying social cognition: a Hebbian perspective. Trends Cogn Sci 8:501–7. doi:10.1016/j.tics.2004.09.005

Khanna P, Carmena JM. 2015. Neural oscillations: beta band activity across motor networks. Curr Opin Neurobiol 32:60–67. doi:10.1016/j.conb.2014.11.010

Kilner JM, Frith CD. 2008. Action observation: inferring intentions without mirror neurons. Curr Biol CB 18:R32–3. doi:10.1016/j.cub.2007.11.008

Kopell N, Ermentrout GB, Whittington MA, Traub RD. 2000. Gamma rhythms and beta rhythms have different synchronization properties. Proc Natl Acad Sci U S A 97:1867–1872. doi:10.1073/pnas.97.4.1867

Lerner Y, Honey CJ, Silbert LJ, Hasson U. 2011. Topographic mapping of a hierarchy of temporal receptive windows using a narrated story. J Neurosci Off J Soc Neurosci 31:2906–15. doi:10.1523/JNEUROSCI.3684-10.2011

Maranesi M, Livi A, Bonini L. 2017. Spatial and viewpoint selectivity for others’ observed actions in monkey ventral premotor mirror neurons. Sci Rep 7:8231. doi:10.1038/s41598-017-08956-1

Mooshagian E, Holmes CD, Snyder LH. 2021. Local field potentials in the parietal reach region reveal mechanisms of bimanual coordination. Nat Commun 12:2514. doi:10.1038/s41467-021-22701-3

Moreno I, de Vega M, León I. 2013. Understanding action language modulates oscillatory mu and beta rhythms in the same way as observing actions. Brain Cogn 82:236–242. doi:10.1016/j.bandc.2013.04.010

Muthukumaraswamy SD, Johnson BW. 2004. Primary motor cortex activation during action observation revealed by wavelet analysis of the EEG. Clin Neurophysiol 115:1760–1766. doi:10.1016/j.clinph.2004.03.004

Nolte G, Ziehe A, Nikulin VV, Schlögl A, Krämer N, Brismar T, Müller K-R. 2008. Robustly Estimating the Flow Direction of Information in Complex Physical Systems. Phys Rev Lett 100:234101. doi:10.1103/PhysRevLett.100.234101

Oostenveld R, Fries P, Maris E, Schoffelen J-M. 2011. FieldTrip: Open source software for advanced analysis of MEG, EEG, and invasive electrophysiological data. Comput Intell Neurosci 2011:156869. doi:10.1155/2011/156869

Qin C, Tan Z, Pan Y, Li Y, Wang Lin, Ren L, Zhou W, Wang Liang. 2017. Automatic and Precise Localization and Cortical Labeling of Subdural and Depth Intracranial Electrodes. Front Neuroinformatics 11. doi:10.3389/fninf.2017.00010

Rao HM, Mayo JP, Sommer MA. 2016. Circuits for presaccadic visual remapping. J Neurophysiol 116:2624–2636. doi:10.1152/jn.00182.2016

Ray S, Maunsell JHR. 2011. Different origins of gamma rhythm and high-gamma activity in macaque visual cortex. PLoS Biol 9:e1000610. doi:10.1371/journal.pbio.1000610

Richter CG, Thompson WH, Bosman CA, Fries P. 2017. Top-Down Beta Enhances Bottom-Up Gamma. J Neurosci 37:6698–6711. doi:10.1523/JNEUROSCI.3771-16.2017

Rizzolatti G, Sinigaglia C. 2016. The mirror mechanism: a basic principle of brain function. Nat Rev Neurosci 17:757–765. doi:10.1038/nrn.2016.135

Simon S, Mukamel R. 2016. Power modulation of electroencephalogram mu and beta frequency depends on perceived level of observed actions. Brain Behav 6. doi:10.1002/brb3.494

Stadler W, Ott DVM, Springer A, Schubotz RI, Schütz-Bosbach S, Prinz W. 2012. Repetitive TMS Suggests a Role of the Human Dorsal Premotor Cortex in Action Prediction. Front Hum Neurosci 6. doi:10.3389/fnhum.2012.00020

Thomas RM, De Sanctis T, Gazzola V, Keysers C. 2018. Where and how our brain represents the temporal structure of observed action. NeuroImage 183:677–697. doi:10.1016/j.neuroimage.2018.08.056

Tia B, Takemi M, Kosugi A, Castagnola E, Ansaldo A, Nakamura T, Ricci D, Ushiba J, Fadiga L, Iriki A. 2017. Cortical control of object-specific grasp relies on adjustments of both activity and effective connectivity: a common marmoset study. J Physiol 595:7203–7221. doi:10.1113/JP274629

Umiltà MA, Kohler E, Gallese V, Fogassi L, Fadiga L, Keysers C, Rizzolatti G, Umilta MA, Kohler E, Gallese V, Fogassi L, Fadiga L, Keysers C, Rizzolatti G. 2001. I know what you are doing. a neurophysiological study. Neuron 31:155–165. doi:S0896-6273(01)00337-3 [pii]

Urgen BA, Saygin AP. 2020. Predictive processing account of action perception: Evidence from effective connectivity in the action observation network. Cortex 128:132–142. doi:10.1016/j.cortex.2020.03.014

van Kerkoerle T, Self MW, Dagnino B, Gariel-Mathis M-A, Poort J, van der Togt C, Roelfsema PR. 2014. Alpha and gamma oscillations characterize feedback and feedforward processing in monkey visual cortex. Proc Natl Acad Sci 111:14332–14341. doi:10.1073/pnas.1402773111

Young CK, Ruan M, McNaughton N. 2017. A Critical Assessment of Directed Connectivity Estimates with Artificially Imposed Causality in the Supramammillary-Septo-Hippocampal Circuit. Front Syst Neurosci 11:72. doi:10.3389/fnsys.2017.00072

Ziehe A, Krämer N, Popescu F, Müller K-R. 2010. Comparison of Granger Causality and Phase Slope Index. J Mach Learn Res - Proc Track 6:267–276.

Zirnsak M, Moore T. 2014. Saccades and shifting receptive fields: anticipating consequences or selecting targets? Trends Cogn Sci 18:621–628. doi:10.1016/j.tics.2014.10.002

