## Supplemental Figure 1 for "Predictive coding during action observation revealed by human electrocorticographic activity"

### Supplementary Materials

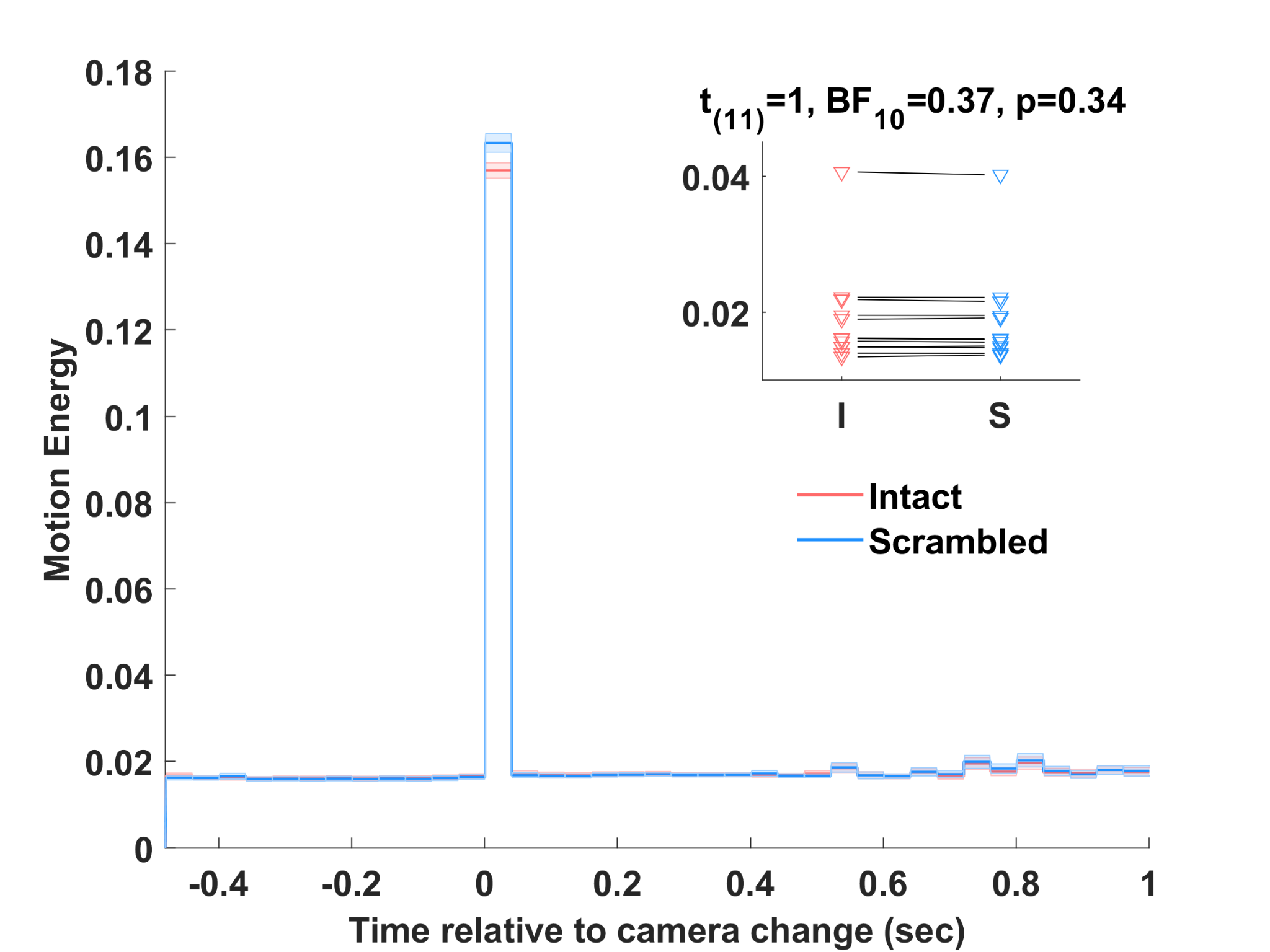

**Figure S1: Motion energy at the transition between acts.** To verify that our use of camera changes ensured that the scrambled sequences do not have significantly more motion energy than our intact movies, motion energy was calculated across consecutive frames as the sum, over all pixels *i*, of the cartesian distance between the vector of (*r,g,b*) values at frame *t* and *t+1*, i.e. $ME=\sum_{i} \sqrt{(r_{t}-r_{t-1})^{2}+(g_{t}-g_{t-1})^{2}+(b_{t}-b_{t-1})^{2}{}}$ , relative to the transition between two acts (*t*=0). Error shading represents the s.e.m. across the 397 transitions over the 12 movies of each condition. As expected, motion energy peaks just following *t*=0, when the camera angle changes, with a trend for 4% more motion energy for scrambled sequences, but the difference does not survive correction for multiple comparison when applying the same statistics as used in Fig 3, 4 and 6 (t-test, corrected for multiple comparison using fdr, q=0.85), and is therefore unlikely to account for the observed differences in the ECoG signal. As some of our results are obtained using the entire movies, as in Figure 2, we also compared motion energy over the entire movies (inset), and again found no significant difference across our conditions when using a matched pair t-test (each triangle represents one of the 12 movies used).
