## Supplemental Figure 2 for "Predictive coding during action observation revealed by human electrocorticographic activity"

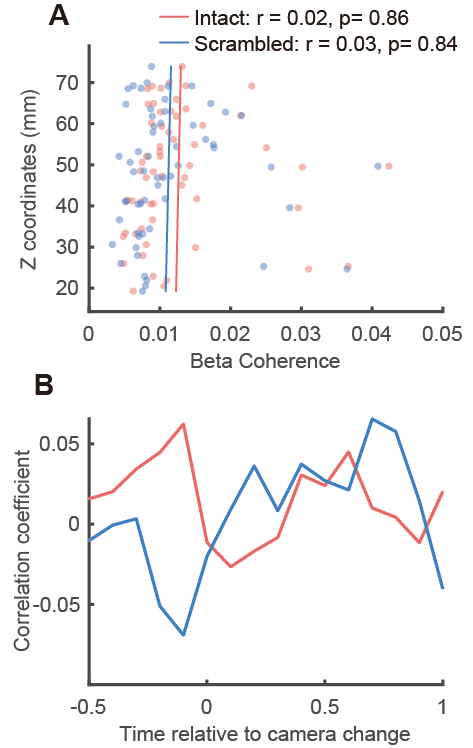

**Figure S2: Location specificity of precentral electrodes in coherence with supramarginal electrodes and its temporal dynamics.**

(A) The beta coherence of precentral and supramarginal electrodes estimated in 0–1s time window relative to camera change, each dot represents a precentral electrode with the location on the Z axis and the average coherence with all supramarginal electrodes. Here we didn't find a correlation between the location of precentral electrodes and the coherence in both conditions. (B) The same correlation estimated in sliding time windows from -0.5 to 1 relative to camera change, no significant correlation was found within this period in both conditions.
