## Supplemental Table 1 for "Predictive coding during action observation revealed by human electrocorticographic activity"

**Table S1: Demographic characteristics of patients.** Bil = bilateral; F = frontal lobe; lat = lateral; Lt = left; med = medial; O = occipital lobe; P = parietal lobe; Rt = right; T = temporal lobe.

| **No** | **Age (y)/Sex** | **Electrode coverage** | **Epileptic foci** | **Etiology and MRI findings** |
| --- | --- | --- | --- | --- |
| 1 | 37/M | Rt TO, Lt T | Rt lat & med T | No lesion |
| 2 | 34/M | bil T | bil T | No lesion |
| 3 | 23/M | Lt TPO | Lt TP | Cavernous hemangioma |
| 4 | 21/F | bil FT | Lt lat T | Ganglioglioma |
| 5 | 39/F | Lt FTP, Rt T | bil med T | No lesion |
| 6 | 30/M | Lt FTP | Lt lat T | No lesion |
| 7 | 23/F | Lt FTP | Lt lat TP | Focal cortical dysplasia |
| 8 | 26/F | bil FTP | bil lat & med T | Post-encephalitis |
| 9 | 22/M | bil FTP | Lt med T | Post-encephalitis |
| 10 | 18/F | Rt FTP | Rt lat T | Rt T ulegyria |
